# Passive infusion of an S2-Stem broadly neutralizing antibody protects against SARS-CoV-2 infection and lower airway inflammation in rhesus macaques

**DOI:** 10.1101/2024.07.30.605768

**Authors:** Christopher T. Edwards, Kirti A. Karunakaran, Elijah Garcia, Nathan Beutler, Matthew Gagne, Nadia Golden, Hadj Aoued, Kathryn L. Pellegrini, Matthew R. Burnett, Christopher Cole Honeycutt, Stacey A. Lapp, Thang Ton, Mark C. Lin, Amanda Metz, Andrei Bombin, Kelly Goff, Sarah E. Scheuermann, Amelia Wilkes, Jennifer S. Wood, Stephanie Ehnert, Stacey Weissman, Elizabeth H. Curran, Melissa Roy, Evan Dessasau, Mirko Paiardini, Amit A. Upadhyay, Ian Moore, Nicholas J. Maness, Daniel C. Douek, Anne Piantadosi, Raiees Andrabi, Thomas R. Rogers, Dennis R. Burton, Steven E. Bosinger

**Affiliations:** Division of Microbiology and Immunology, Emory National Primate Research Center, Emory University, Atlanta, GA 30329, USA; Department of Pathology, Microbiology & Immunology, Vanderbilt University, Nashville, TN 37235, USA; Department of Immunology and Microbiology, The Scripps Research Institute, La Jolla, CA 92037, USA; Mayo Clinic Medical Scientist Training Program, Mayo Clinic College of Medicine and Science, 200 First Street SW, Rochester, Minnesota 55356, USA; Vaccine Research Center; National Institute of Allergy and Infectious Diseases, National Institutes of Health, Bethesda, MD, USA; Tulane National Primate Research Center, Covington, LA, USA; Emory National Primate Research Center Genomics Core, Emory National Primate Research Center, Emory University, Atlanta, GA 30329, USA; Division of Infectious Diseases, Department of Medicine, Emory University School of Medicine, Atlanta, GA, USA; Division of Animal Resources, Emory National Primate Research Center, Emory University, Atlanta, GA 30329, USA; Division of Pathology, Emory National Primate Research Center, Emory University, Atlanta, GA 30329, USA; Division of Histology, Emory National Primate Research Center, Emory University, Atlanta, GA 30329, USA; Emory Vaccine Center, Emory National Primate Research Center, Atlanta, Georgia, USA; Department of Pathology and Laboratory Medicine, Emory University School of Medicine, Atlanta, GA, USA; IAVI Neutralizing Antibody Center, The Scripps Research Institute, La Jolla, CA 92037, USA; Consortium for HIV/AIDS Vaccine Development (CHAVD), The Scripps Research Institute, La Jolla, CA 92037, USA; Division of Infectious Diseases, Department of Medicine, University of California, San Diego, La Jolla, CA 92037, USA; Ragon Institute of Massachusetts General Hospital, Massachusetts Institute of Technology, and Harvard University, Cambridge, MA 02139, USA

## Abstract

The continued evolution of SARS-CoV-2 variants capable of subverting vaccine and infection-induced immunity suggests the advantage of a broadly protective vaccine against betacoronaviruses (β-CoVs). Recent studies have isolated monoclonal antibodies (mAbs) from SARS-CoV-2 recovered-vaccinated donors capable of neutralizing many variants of SARS-CoV-2 and other β-CoVs. Many of these mAbs target the conserved S2 stem region of the SARS-CoV-2 spike protein, rather the receptor binding domain contained within S1 primarily targeted by current SARS-CoV-2 vaccines. One of these S2-directed mAbs, CC40.8, has demonstrated protective efficacy in small animal models against SARS-CoV-2 challenge. As the next step in the pre-clinical testing of S2-directed antibodies as a strategy to protect from SARS-CoV-2 infection, we evaluated the *in vivo* efficacy of CC40.8 in a clinically relevant non-human primate model by conducting passive antibody transfer to rhesus macaques (RM) followed by SARS-CoV-2 challenge. CC40.8 mAb was intravenously infused at 10mg/kg, 1mg/kg, or 0.1 mg/kg into groups (n=6) of RM, alongside one group that received a control antibody (PGT121). Viral loads in the lower airway were significantly reduced in animals receiving higher doses of CC40.8. We observed a significant reduction in inflammatory cytokines and macrophages within the lower airway of animals infused with 10mg/kg and 1mg/kg doses of CC40.8. Viral genome sequencing demonstrated a lack of escape mutations in the CC40.8 epitope. Collectively, these data demonstrate the protective efficiency of broadly neutralizing S2-targeting antibodies against SARS-CoV-2 infection within the lower airway while providing critical preclinical work necessary for the development of pan–β-CoV vaccines.

**AUTHOR SUMMARY:** In this study, we explore the development of a broadly protective vaccine against betacoronaviruses (β-CoVs), including SARS-CoV-2. We focused on monoclonal antibodies (mAbs) from individuals who recovered-vaccinated donors capable of neutralizing many variants of SARS-CoV-2 and other β-CoVs. Unlike current vaccines that target the S1 region of the virus, these mAbs target a highly conserved S2 region of the spike protein. One antibody, CC40.8, showed promising results in small animal models. To further test its effectiveness, we infused CC40.8 into rhesus macaques at different doses and then challenged them with SARS-CoV-2. We found that higher doses of CC40.8 significantly reduced viral loads and inflammation in the lower airway. Additionally, there were no escape mutations in the targeted region, suggesting that the virus could not easily evade the antibody. Our findings highlight the potential of S2-targeting antibodies to protect against SARS-CoV-2 and support the development of vaccines that can broadly protect against various β-CoVs.

*Conflicting Interests:* RA, TFR, and DRB are listed as inventors on pending patent applications describing the SARS-CoV-2 and HCoV-HKU1 S cross-reactive antibodies. DRB and RA are listed as inventors on a pending patent application describing the S2 stem epitope immunogens identified in this study. DRB is a consultant for IAVI. All other authors declare that they have no competing interests.

**ONE SENTENCE SUMMARY:** Pan-beta-coronavirus neutralizing mAb CC40.8 reduces SARS-CoV-2 viral loads and inflammation within the lower airway of infected rhesus macaques and provides pre-clinical support for S2-directed immunization strategies.

## INTRODUCTION

Since its emergence in late 2019, severe acute respiratory syndrome coronavirus 2 (SARS-CoV-2) has led to over 700 million cases of coronavirus disease 2019 (COVID-19), resulting in over 6 million deaths(1). While alphacoronaviruses HCoV-229E and HCoV-NL63, and betacoronaviruses (β-CoVs) HCoV-OC43 and HCoV-HKU1 are endemic to humans, typically causing mild disease, they still pose a serious threat to at-risk populations such as the elderly and immunocompromised (2-4). β-CoVs SARS-CoV-1 (severe acute respiratory syndrome coronavirus 1) and MERS-CoV (Middle East respiratory syndrome CoV) both arose from zoonotic transmission events within the last 20 years and are associated with high morbidity and mortality in humans (4-6). Together with the COVID-19 pandemic, these transmission events and subsequent public health crises highlight the urgent need for proactive measures to prevent a future coronavirus epidemic.

Today, vaccination remains the most utilized prophylactic strategy against severe COVID-19, with over 13.5 billion doses of SARS-CoV-2 vaccines across various platforms administered worldwide (7). The majority of approved SARS-CoV-2 vaccines seek to induce neutralizing antibodies against the surface spike (S) glycoprotein, particularly the receptor binding domain (RBD) regardless of platform (8-20). Due to their highly efficient immune responses, rapid development, and ease of scalability, messenger ribonucleic acid (mRNA)-based vaccines developed separately by Moderna (mRNA-1273) and Pfizer/BioNTech (BNT162b2) were the first to be approved by the US Food and Drug Administration and European Medicines Agency, both encoding a prefusion-stabilized, full-length S protein (8, 21-23). Approval and administration of viral vector and protein subunit-based vaccines also utilizing full-length S proteins have since followed (24, 25). Even though full-length S immunogens do include both the S1 and S2 subunits, the majority of IgG responses target the highly immunogenic S1(26, 27).

The majority of human coronavirus (HCoV) infections elicit strain-specific neutralizing antibody responses (28, 29). Only 10-13% of convalescent COVID-19 donors exhibit some degree of neutralizing capacity against multiple β-CoVs (30-32). Several theories have been proposed to explain the rarity of broad neutralizing humoral immunity against β-CoVs, including “antigenic sin” driven by previous coronavirus infection towards non-neutralizing or variant-specific epitopes, steric difficulty in targeting conserved epitopes, and disfavored somatic hypermutation pathways (33). Broad neutralizing responses have been elicited via heterologous coronavirus (CoV) S subunit vaccinations in human immunoglobulin (Ig) locus transgenic mice. However, much more remains to be investigated on conserved epitope targeting antibody development in more clinically relevant models (34, 35).

SARS-CoV-2 has continued to evolve to escape immune pressures applied by B and T cell memory conferred by both infection and vaccination. During the first two years of the pandemic, different lineages designated as “variants of concern” (VOCs) by the World Health Organization (WHO) have repeatedly emerged from distinct temporal and geographic landscapes. These VOCs were defined by up to 16 point mutations and a deletion of 7 nucleotides that conferred an overall fitness advantage over co-circulating variants (36-40). However, after the omicron variant emerged in November 2021, its sub-lineages quickly outcompeted other variants and ushered in what was described as a “fourth-wave” of the pandemic (41). Harboring a variety of mutations within the S protein, including 30 amino acid substitutions, a deletion of 6 amino acids, and an insertion of 3 new amino acids, the omicron VOCs represented a phylogenetically distant lineage compared to previous VOCs (36, 42). Omicron and its sub-lineages are characterized by their high transmissibility, less severe disease, and resistance against both previously approved therapeutic antibodies and those from convalescent patients or vaccinated individuals (43-45). These characteristics are largely attributed to the concentration of over 15 amino acid substitutions within the ACE2 RBD and 9 substitutions within the S1 NTD. The swift rise of a phylogenetically distinct variant able to circumvent existing intervention strategies underscores the prudence of reducing reliance on S1-targeted humoral immunity in favor of a more universal CoV prevention approach.

In terms of S1-targeted immunity, the majority of neutralizing antibodies generated against SARS-CoV-2 target the highly immunogenic RBD contained within the S1 subunit (46, 47). The RBD is responsible for engaging the human angiotensin-converting enzyme 2 (hACE2) on the surface of host cells within the airways (46-50). While the RBD is the major target of neutralizing antibodies, this region exhibits considerable variation between HCoVs and most RBD-neutralizing epitopes are likely to be susceptible to antigenic drift (2, 51-53). The S2 subunit, responsible for mediating membrane fusion, is more conserved among β-CoVs, with both the stem helix region and fusion peptide-containing epitopes targeted by broadly neutralizing antibodies (bnAbs) (35, 54, 55). Antibodies targeting this region typically exhibit lower neutralizing potency than those targeting the RBD, but have been shown to retain similar protective efficacy in vivo, possibly due to Fc-mediated effector mechanisms (30). Recently, several S2-targeting antibodies with neutralizing breadth against multiple SARS-CoV-2 VOCs or across β-CoVs have been described by us and others (55). We have shown that these antibodies target a conserved 25 aa stem-helix region in the S2 domain, use a remarkably restricted set of V genes, and were isolated most frequently in subjects with hybrid immunity (natural infection followed by vaccination) but rarely in individuals exposed to either SARS-CoV-2 by infection or vaccination alone (41).

We have previously shown that CC40.8, a monoclonal antibody (mAb) isolated from a peripheral blood mononuclear cell (PBMC) sample of a 62-year-old convalescent donor, exhibits broad reactivity against β-CoVs by targeting the conserved stem helix (SH) epitope of the S2 region (56). In addition to neutralizing both SARS-CoV-1 and VOCs of SARS-CoV-2, CC40.8 protects against weight loss and reduces viral burden in SARS-CoV-2 challenge *in vivo* when passively infused into hACE2 mice and Syrian hamsters (30). However, the efficacy of S2-directed humoral immunity at preventing COVID-19 pathology has yet to be described in non-human primates (NHPs). Owing to the genetic and physiological similarities to humans, NHPs are a highly relevant model for investigating both the pathology of SARS-COV-2 infection, as well as the efficacy of vaccination and therapeutic strategies against the virus (57). NHPs, including rhesus macaques (RMs), have an ACE2 receptor that is nearly identical to humans, proving valuable when testing vaccine-mediated protection against different variants (58-62). Using β-CoV bnAb CC40.8 in the mild-moderate disease model of SARS-CoV-2–infected RMs, our data demonstrate the ability of S2-mediated neutralizing antibodies to limit viral load and inflammation within the lower airway.

## RESULTS

### CC40.8 reduces SARS-CoV-2 replication in the lower airway of rhesus macaques

We have previously shown that β-CoV bnAb CC40.8 consistently neutralizes SARS-CoV-2 VOCs *in vitro* and significantly protects from weight loss and lowers airway viral loads *in vivo* in small animal models(30). To determine the *in vivo* efficacy of CC40.8 in the non-human primate model, we conducted passive antibody transfer followed by SARS-CoV-2 challenge. CC40.8 mAb was intravenously infused at 10mg/kg, 1mg/kg, 0.1 mg/kg doses into groups of six rhesus macaques (Fig. 1A, Table S1), and an HIV-specific antibody (PGT121, 10mg/kg) negative control was intravenously administered to a control group (n=6). All four groups were challenged with SARS-CoV-2 (2019-nCoV/USA-WA1/2020) intranasally and intratracheally with a combined total of 1.1 x 10^6^ plaque-forming units (PFU) five days following antibody infusion. Three animals per experimental group were euthanized at 7 and 8 days post infection (dpi) each. Animals were scored according to the Coronavirus Vaccine and Treatment Evaluation Network standard clinical assessment at cage-side (Table S2) and during anesthetic (Table S3) accesses. No differences were observed in cage-side, anesthetic, or total clinical scores between treatment groups. No treatment group experienced weight changes (Fig. S2C), and vitals including rectal temperature, respiratory rate, heart rate, and SpO2 also did not differ between experimental groups (Fig S2B-E). Levels of CC40.8 and PGT121 in the sera and bronchial alveolar lavages (BALs) accurately reflected experimental dosage across all animals from -4 dpi through 7/8 dpi (Fig 1D-1E).

**Fig. 1.**
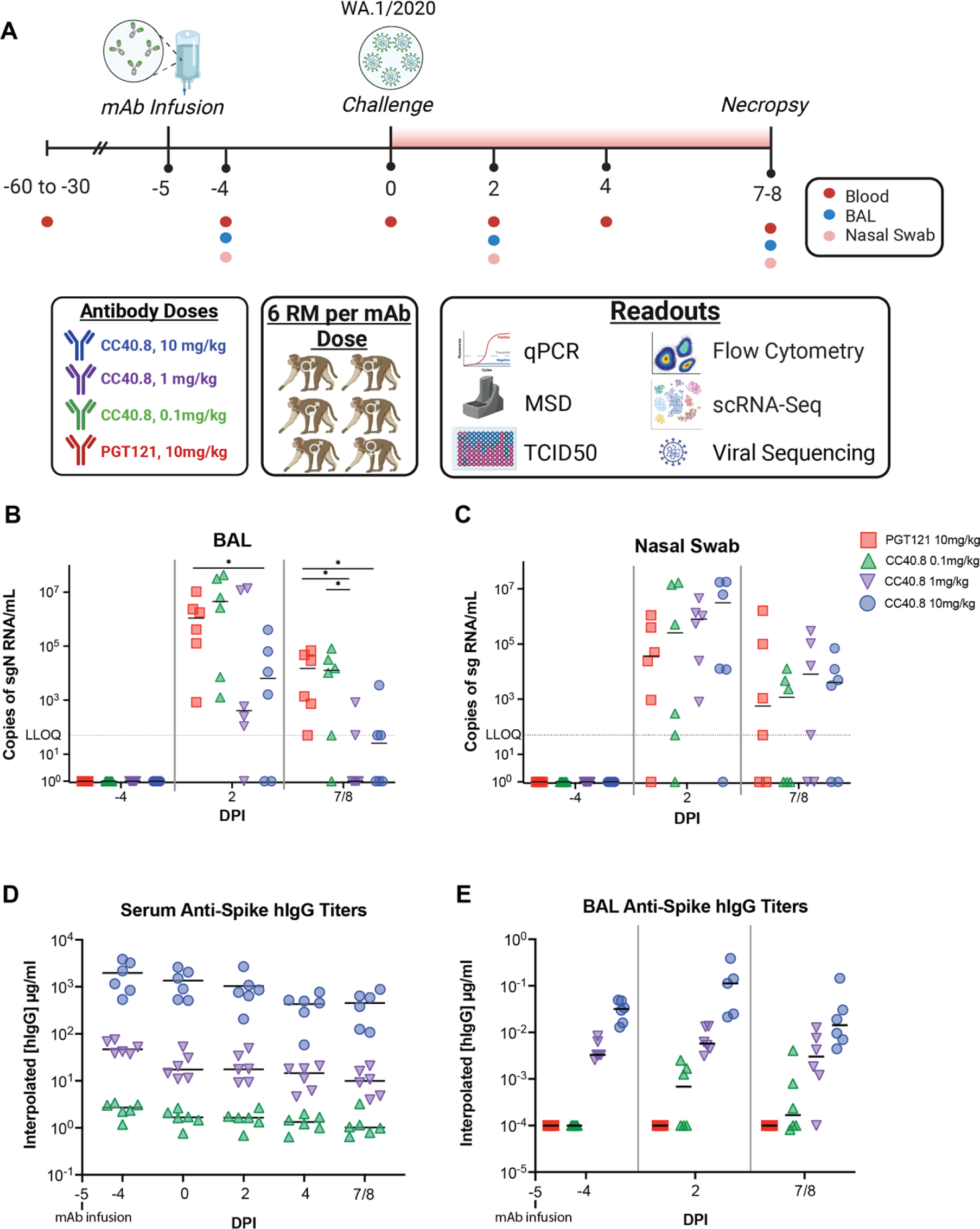
Preinfusion of S2-targeting bnAb CC40.8 reduced viral loads in RMs on SARS CoV-2 challenge. (**A**) 24 RMs (6 females and 18 males; mean age of 5 years and 11 months old; range 5-6 years old) were infused intravenously 5 days pre infection with either a 0.1 mg/kg, 1mg/kg, or 10mg/kg concentration of SARS-CoV-2 bnAb CC40.8 or with control mAb PGT121, with each experimental group consisting of 6 RMs (1 female and 5 males). RMs were screened for preexisting, SARS-CoV-2 spike specific antibodies prior to CC40.8 administration RMs were euthanized at 7 dpi (*n* = 3 RMs per treatment arm) or 8 dpi (*n* = 3 RMs per treatment arm). Levels of SARS-CoV-2 sgRNA N in BAL (**B**) and nasopharyngeal swabs (**C**). (**D**) Anti-Spike hIgG titers in the serum (**D**) and BAL (**E**) measured via ELISA. Control PGT121-treated RMs are depicted with red squares, CC40.8 0.1mg/kg-treated RMs depicted with green upward pointing triangles, CC40.8 1mg/kg-treated RMs depicted with purple downward pointing triangles, CC40.8 10mg/kg-treated RMs depicted with blue circles. Black lines represent the median viral loads for each treatment group at each time point. Statistical analyses were performed using nonparametric Mann-Whitney tests. \**P* < 0.05.

Viral titers were determined via quantitative polymerase chain reaction (qPCR) for both subgenomic RNA (sgRNA) and genomic RNA (gRNA) to measure replicating virus and to verify inoculation, respectively (Fig. 1B-C, S1A-B). BAL and nasal swab sgRNA levels were reproduced by an independent laboratory and further confirmed by gRNA quantification (fig. S1E-F). 10mg/kg CC40.8-treated animals exhibited significant reductions (p = 0.039) in SARS-CoV-2 subgenomic N (sgN) levels within the BAL at 2 dpi compared to both negative control (PGT121 10mg/kg) and 0.1 mg/kg dose treated animals (Fig. 1B). In addition, BAL subgenomic E (sgE) viral titers at 2 dpi showed significant reduction between the 10 mg/kg treated animals compared to the negative control (p = 0.042) and 0.1 mg/kg dose (p = 0.042) (Fig. 1B). Levels of replication-competent virus within the BAL at 2 dpi trended similarly (Fig S1). At 7/8 dpi, the majority of CC40.8 1 mg/kg and 10 mg/kg treated animals exhibited sgN loads at or below the lower level of quantification, while negative control and 0.1 mg/kg dosed animals retained above 100,000 copies of sgN per mL (Fig. 1B, Fig S1E). Compared to the negative control, 7/8 dpi sgN viral loads within the BAL of 1.0 mg/kg and 10 mg/kg treatment groups were significantly reduced (p = 0.015 and p = 0.011 respectively), and a significant difference was also observed between 0.1 mg/kg treatment and 1.0 mg/kg treatment (p=0.045) (Fig. 1B). Despite these observations within the BAL and lung tissues, no significant differences in sgN or sgE viral titers were observed within the upper airway in any of the RM groups (Fig. 1C, Figs. S1F, S1H, S1J).

### CC40.8 reduces lower airway infiltration of inflammatory monocyte and macrophage populations during SARS-CoV-2 infection

Several studies have reported perturbed macrophage populations within the airways of rhesus macaques during SARS-CoV-2 infection(63-65). In our prior work in the RM model, we identified the predominant macrophage subsets producing inflammatory cytokines in the lower airway following SARS-CoV-2 infection using single-cell RNA sequencing (scRNA-Seq) as CD163+MRC1+TREM2+ and CD163+MRC1− macrophages(61, 66), and that blocking the recruitment of these subsets abrogated associated inflammatory signaling(61, 66). To expand on our characterization of these subsets, we developed a multi-parametric flow cytometry panel to assess changes in frequency within the BAL of alveolar macrophage populations (CD163+ MRC1+) and of non-tissue-resident macrophage populations (CD163+ MRC1-) (Fig. 2A Fig. S3A). Treatment with 1.0 mg/kg and 10 mg/kg of CC40.8 resulted in significant reductions (p = .008658 and p = .030303, respectively) in infiltrating CD163+ MRC1-macrophage populations within the BAL at 2 dpi (Fig 2B and 2E). Similarly, 1.0 mg/kg and 10 mg/kg groups maintained their CD163+ MRC1+ alveolar macrophage populations across the course of infection, while the 0.1 mg/kg dose and control groups exhibited significant reductions in this population compared to 10 mg/kg treated animals at 2 dpi (p = .007937 and p = .008658 respectively) (Fig. 2C and 2F).

**Fig. 2.**
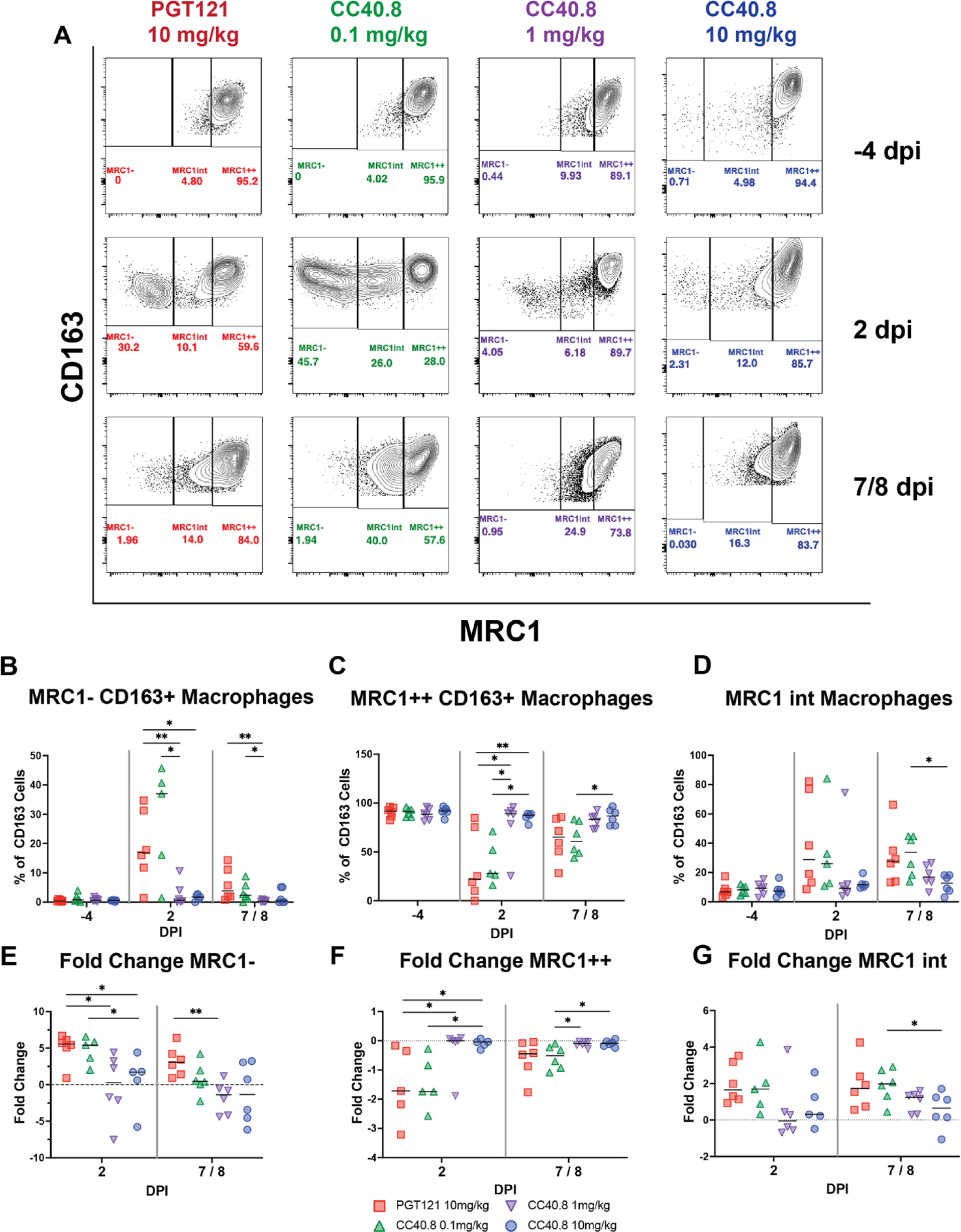
CC40.8-treated RMs had lower frequencies of inflammatory CD163+ MRC1-macrophages compared with PGT121-treated RMs. (**A**) Representative staining of macrophages for CD163 and MRC1 in the BAL at -4, 2, and 7/8 dpi with frequency as a percentage of total CD163+ cells. Macrophages were gated on singlets, CD45+, FSC and SSC characteristic of granulocytes and alveolar macrophages, live cells, and CD14+ populations. (**B** to **D**) Frequency as a percentage of total CD163+ cells for (**B**) MRC1-, (**C**) MRC1 intermediate and (**D**) MRC1++ cells. (**E** to **F**) Fold change from -4 dpi baseline as a percentage of total CD163+ cells for (**E**) MRC1-, (**F**) MRC1 intermediate, and (**G**) MRC1++ cells. Control PGT121-treated RMs are depicted with red squares, CC40.8 0.1mg/kg-treated RMs depicted with green upward pointing triangles, CC40.8 1mg/kg-treated RMs depicted with purple downward pointing triangles, CC40.8 10mg/kg-treated RMs depicted with blue circles. Black lines represent the median frequency or fold change in RMs from each respective treatment group. Statistical analyses were performed using nonparametric Mann-Whitney tests. \**P* < 0.05, \*\**P* < 0.01

We also quantified the impact of CC40.8 treatment on airway macrophage populations using droplet based scRNA-Seq. Our previous work delineated the major subsets of lung macrophages driving inflammatory and anti-inflammatory cytokine production within the alveolar space during SARS-CoV-2 infection(61, 66). We employed the same approach in this study, using previously generated scRNA-Seq data from uninfected RMs as a reference to map and annotate 107,830 cells captured from the BAL from all 24 animals (Fig 3A, Fig. S5A-C). Consistent with prior studies, the vast majority of annotated cells were macrophages and monocytes, and transcriptomic analysis identified four major macrophage/monocyte subsets: (i) CD163+MRC1+ resident alveolar macrophages; (ii) macrophages similar to infiltrating monocytes expressing CD163+MRC1+TREM2 (iii) CD163+ MRC1-interstitial-like macrophages; and (iii) CD16+ monocytes (Fig. 3B) (66, 67). scRNA-Seq demonstrated that animals infused with 10 mg/kg of CC40.8 had near complete abrogation of the influx of CD163+MRC1-macrophages to the lower airway at 2 dpi compared to control animals (p = .030303) (Fig. 3C0.1 mg/kg treated animals had significantly higher frequencies of these interstitial-like macrophages at both 2 dpi and 7 / 8 dpi compared to 10 mg/kg treated animals (p = .031746 and p = .015152 respectively), consistent with our flow cytometry data (Fig. 3C). The frequency of CD163+MRC1-macrophages at 2 dpi within the BAL significantly correlated with BAL viral load (Fig. S6E). While we did not measure significant differences between experimental groups in CD163+MRC1+ alveolar macrophage frequency via scRNA-Seq, there was a trend towards a CC40.8 dose response in maintaining this population in the airway (Fig 3E). We also detected a significant reduction in CD16+ monocyte populations within the BAL of 1.0 mg/kg and 10 mg/kg treatment groups compared to controls (p = .021645 and p = .028139 respectively) (Fig. 3F). Interestingly, we did not measure significant differences between treatment conditions in the frequency of CD163+MRC1+TREM2+ macrophages, however, this may be due to timing of sampling, as our prior work has demonstrated that this subset peaks at 4 dpi (Fig. 3D)(66). Collectively these data demonstrate that S2 targeted neutralization of SARS-CoV-2 by CC40.8 is capable of eliminating the recruitment of inflammatory myeloid cells into the lower airway during infection.

**Fig. 3.**
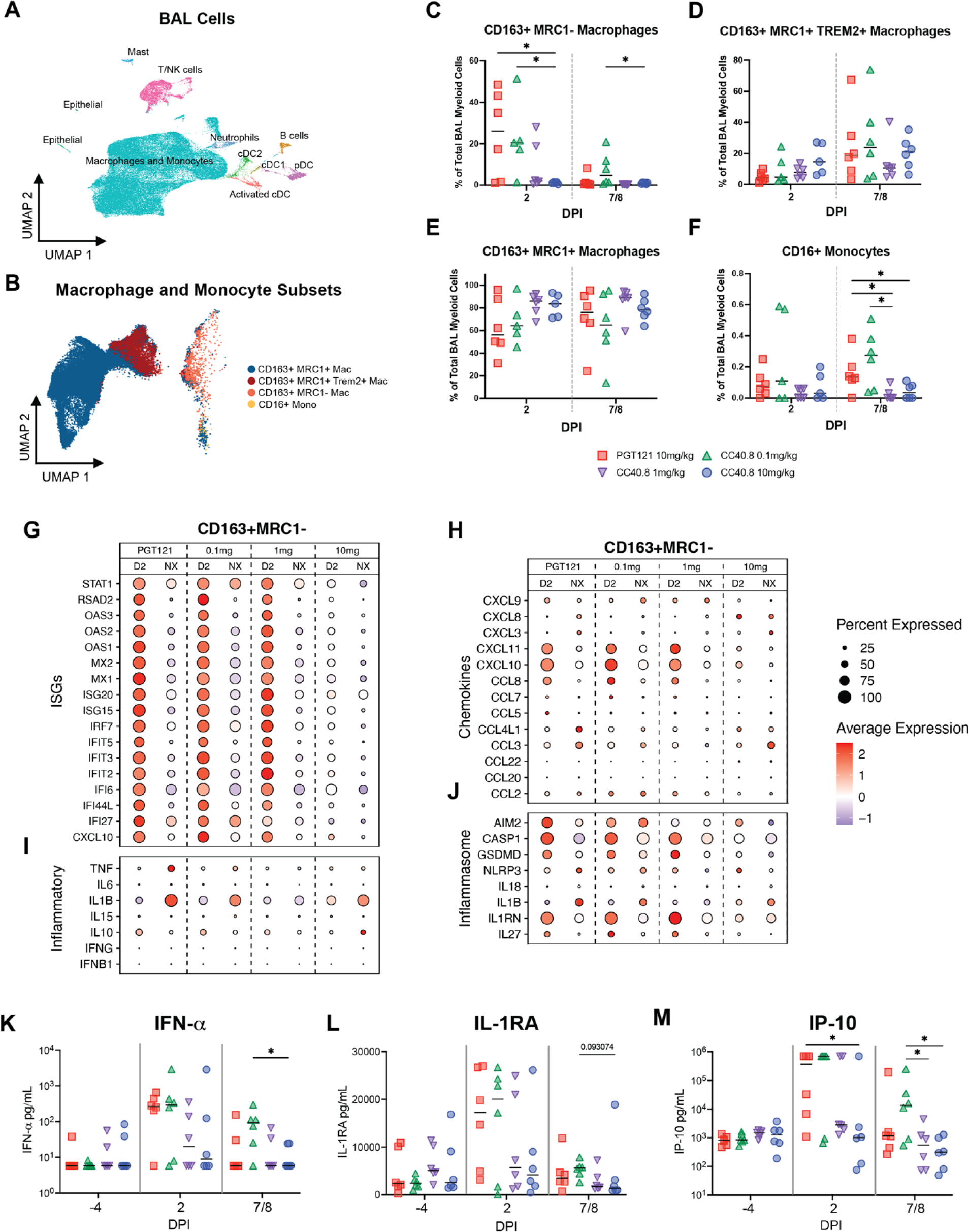
Effect of CC40.8 treatment on gene expression of BAL single cells during SARS-CoV-2 infection using 10X. **A**) Uniform Manifold Approximation and Projection (UMAP) of BAL samples (107830 cells) integrated using reciprocal principal components analysis (PCA) showing cell type annotations. Captures were performed on BAL cells from all RMs at 2 and 7/8 dpi. (**B**) Mapping of macrophage/monocyte cells in the BAL of SARS-CoV-2–infected PGT121- and CC40.8-treated RMs to different lung macrophage/monocyte subsets from healthy RMs (67). (**C** to **F**) Percentage of different macrophage/monocyte subsets of all the macrophage/monocytes in BAL at 2 and 7/8 dpi from PGT121- and CC40.8-treated RMs. Frequency as a percentage of total CD163+ cells for (**C**) MRC1-, (**D**) MRC1+ TREM2+, (**E**) MRC1++, and (**F**) CD16+ monocytes. (**G** to **J**) Dot plots showing the expression of selected (**G**) ISGs, (**I**) inflammatory genes, (**H**) chemokines, and (**J**) inflammasome genes in CD163+ MRC1-macrophages. The size of the dot indicates the percentage of cells that express a given gene, and the color indicates the level of expression. (**K** to **M**) Fold change of cytokines and chemokines in BAL fluid relative to −4 dpi measured by MSD immunoassay. Control PGT121-treated RMs are depicted with red squares, CC40.8 0.1mg/kg-treated RMs depicted with green upward pointing triangles, CC40.8 1mg/kg-treated RMs depicted with purple downward pointing triangles, CC40.8 10mg/kg-treated RMs depicted with blue circles. Black lines represent the median frequency or fold change in RMs from each respective treatment group. Statistical analyses were performed using two-sided nonparametric Mann-Whitney tests. \**P* < 0.05.

We also observed significant reductions in protein levels of inflammatory cytokines measured within the BAL of 10 mg/kg CC40.8-treated RM following infection (Fig. 3K-M, Fig S4A-J). While both control and 0.1 mg/kg treated animals trended towards higher levels of IFNα at both 2dpi and 7 / 8 dpi, we observed significant differences at 7 / 8 dpi between the 0.1 mg/kg and 10 mg/kg treated groups (p = .021645) (Fig. 3K). IP-10, previously identified as a biomarker for COVID-19 severity, was significantly reduced in 10 mg/kg treated groups at 2 dpi compared to controls (p = .04329), and in both 1.0 mg/kg and 10 mg/kg groups at 7 / 8 dpi compared to 0.1 mg/kg dose (p = .041126 and p = .025947, respectively) (Fig 3M). G-CSF levels were also significantly elevated in 0.1 mg/kg treated animals at 2 dpi (p = .041126) (Fig.S4A). Many of the measured proinflammatory cytokines and chemokines levels, including TNF-a, IL-1b, and IL-6 correlated highly significantly with BAL SARS-CoV-2 titers (Supplementary Table S4).

### CC40.8 abolishes gene expression programs of inflammation driven by infiltrating macrophages following SARS-CoV-2 infection

To further investigate the impact of CC40.8 treatment on pulmonary macrophages within the alveolar space during early SARS-CoV-2 infection, we identified transcriptional changes in the CD163+MRC1+TREM2+, CD163+ MRC1+ and CD163+ MRC1-macrophage populations. Consistent with the reduction in sgRNA in the BAL, 10 mg/kg treatment of CC40.8 abrogated ISG expression in CD163+ MRC1-, CD163+ MRC1+, and CD163+MRC1+TREM2 macrophages at 2 dpi, with 1.0 mg/kg treated animals also showing reduced ISG expression at this time point (Fig 3G, Fig S6C). Also, consistent with previously published data, the CD163+MRC1− population produced the majority of transcripts for the inflammatory cytokines TNF and IL6, and transcripts for the pro-inflammatory chemokines CCL8, CXCL10, and CXCL11 across experimental groups (Fig 3G-H, Fig S6C). The 10 mg/kg CC40.8 treatment group demonstrated a significant reduction of the levels of these transcripts in the CD163+ MRC1-subset. Interestingly, while we observed broad expression of ISGs in the CD163+ MRC1-macrophages, this treatment group exhibited the highest expression levels of the pro-inflammatory cytokine IL1B at 2 dpi.

Acute SARS-CoV-2 infection results in activated monocytes and macrophages, which undergo inflammasome-mediated pyroptosis (68). This process has been shown to induce secondary inflammation in non-immune cell subsets within the lungs of RM (69). We observed CC40.8-dependent reductions of several inflammasome-associated genes in CD163+ MRC1− macrophages within the BAL (Fig. 3J). In 10 mg/kg treated animals, we observed reductions in the expression of CASP1, GSDMD, IL1RN, IL27 at 2 dpi and AIM2 and NLRP3 at 7/8 dpi compared to control and 0.1 mg/kg treated animals. 1.0 mg/kg dose CC40.8 treatment resulted in reductions in AIM2 at 2 dpi, and AIM2, IL1B, and NLRP at 7/8 dpi. Consistent with our previous work, reductions in inflammasome-associated genes were most prominent in CD163+MRC1− cells, with smaller effects observed in the CD163+MRC1+TREM2 and CD163+MRC1+ populations (Fig. 3J, S6C).

### CC40.8 treatment reduced inflammation within lung during SARS-CoV-2 infection

To further investigate the effect of CC40.8 mAb treatment within the lower airway, scRNA-seq was conducted on cell suspensions prepared from caudal (lower) lung lobe sections obtained at necropsy (7/8 dpi). Our previous work explored SARS-CoV-2 driven inflammation within lung cell subsets [61,66], and we employed a similar approach in this study to annotate 101,766 cells captured from caudal lung tissues from RM treated with PGT121 (n=3), CC40.8 10mg/kg (n=3), and CC40.8 0.1mg/kg (n=3) at 7/8 dpi (Fig 4A, Fig. S7A-D). Cells were classified into four major groups based on the expression of canonical markers (epithelial, lymphoid, myeloid, and “other”) and then each clustered separately (Fig 4A). Subsets within each major group were defined by the expression of marker genes (Fig. S7 A-D). Pathway analysis at 7/8 dpi revealed stark contrast in IFN-I signaling between PGT121 and CC40.8 10mg/kg treated animals (Fig. 4B), consistent with contrasting BAL and lung tissue viral loads measured at the same time point (Fig 1A, Fig S1D). IFN-I related gene sets were observed to be diminished in CC40.8 10mg/kg animals across several cell subsets in the lung, especially when compared to CD163+ MRC1+ macrophages, non-classical monocytes, pDCs, cDC2’s and adventitial fibroblasts in PGT121 treated animals (Fig. 4B). Additionally, analysis of singular genes revealed at least 9 ISGs that were significantly lower in the adventitial fibroblasts, CD163+ MRC1+ macrophages, CD163+ MRC1-macrophages, and neutrophils of CC40.8 10mg/kg treated animals compared to PGT121 controls (Fig. 4C).

**Fig. 4.**
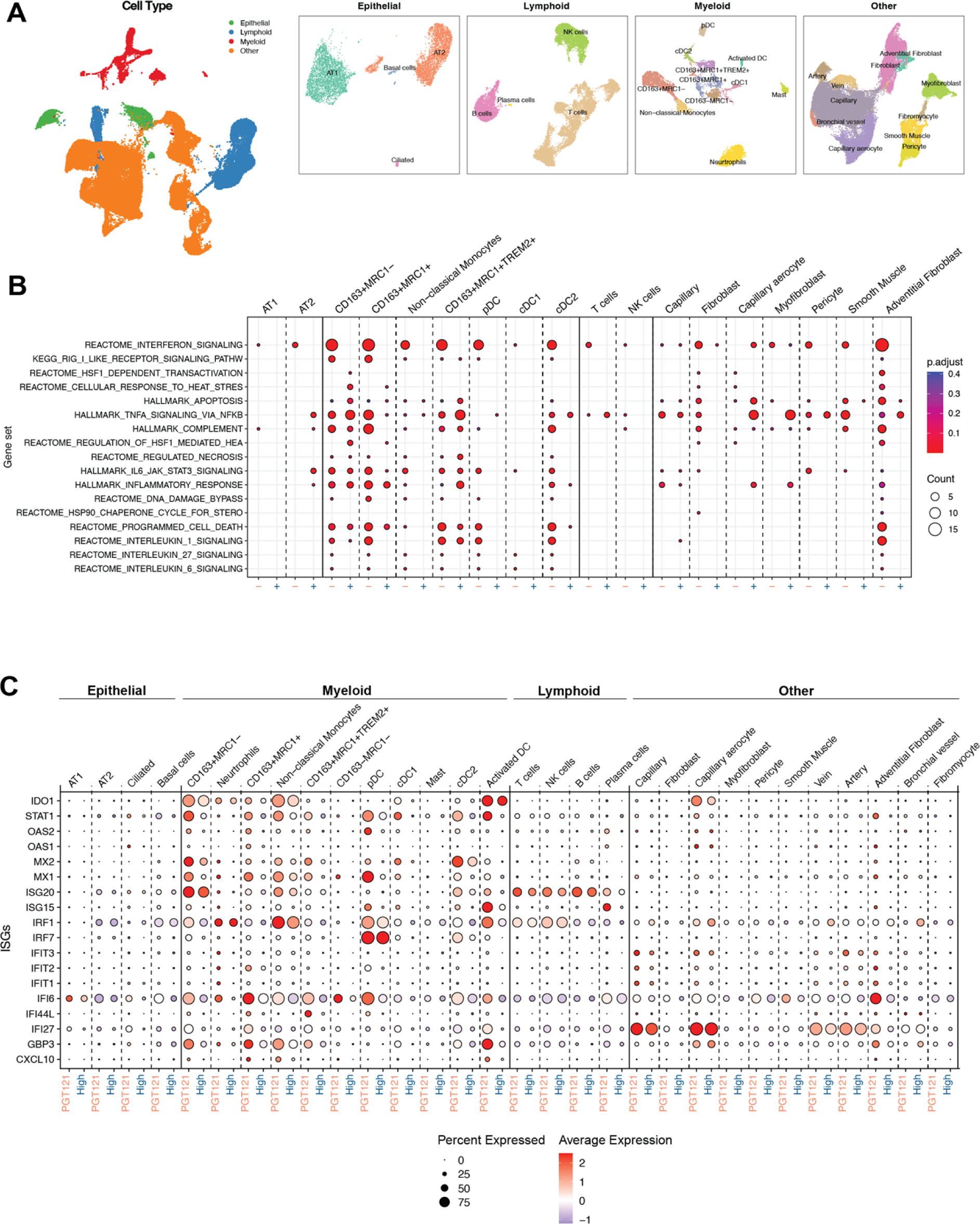
Effect of CC40.8 treatment on lung cells during SARS-CoV-2 infection. (**A**) UMAP based on reciprocal PCA of lung single cells (101,766 cells) collected at 7/8 dpi (*n* = 3 PGT121, 3 CC40.8 0.1mg/kg, and 3 CC40.8 10mg/kg). The cells were classified into four broad categories—epithelial, lymphoid, myeloid, and other, followed by subsetting and separate clustering within each category. UMAPs for each category with cell type annotations are also shown. (**B**) Selected gene sets that were found to be enriched (*P*-adjusted value < 0.05) in lung cells from PGT121-treated RMs compared to CC40.8 10mg/kg-treated RMs at 7/8 dpi based on overrepresentation analysis using Hallmark, Reactome, Kyoto Encyclopedia of Genes and Genomes, and BioCarta gene sets from MSigDB. The size of the dots represents the number of genes that were enriched in the gene set, and the color indicates the *P*-adjusted value. The gene set IDs in order are M983, M15913, M27255, M27253, M5902, M5890, M5921, M27250, M41804, M5897, M5932, M27698, M27251, M29666, M27436, M27895, M27897, and M1014. (**C**) Dot plots showing gene expression in lung cells present at higher frequencies from PGT121- and CC40.8-treated macaques at 7/8 dpi. Plot is organized by epithelial, myeloid, lymphoid and other subsets. The size of the dot represents the percentage of cells expressing a given gene, and the color indicates the average expression.

### CC40.8 treatment did not select for mutations in the S2 stem-helix epitope

To evaluate within-host SARS-CoV-2 evolution and potential antibody escape, we performed full viral genome sequencing using RNA from BAL fluid. We analyzed consensus mutations and intrahost single-nucleotide variants (iSNVs) compared to the sequence of the infecting viral stock. 11 animals yielded sequence data from both the 2 dpi and 7/8 dpi intervals and 10 animals yielded sequence data at solely the 2 dpi timepoint (Data File S8). Three animals (from the CC40.8 1mg/kg or 10 mg/kg groups) had insufficient SARS-CoV-2 RNA for viral genome sequencing.

Very few consensus-level changes were observed across the genome, but we identified numerous iSNVs (Fig. S9). Within-sample diversity, as measured by the average Shannon entropy across the genome within each sample, increased in PGT121-treated control animals but remained constant in CC40.8-treated animals (Fig. 5A). While the low number of animals with paired sequences available hindered robust statistical analysis, there seemed to be a dose dependent decrease in mean entropy for CC40.8 treated groups (Fig. 5B). These results suggest that CC40.8 treatment does not enhance within-host SARS-CoV-2 diversification over the course of infection and may instead limit it.

**Fig. 5.**
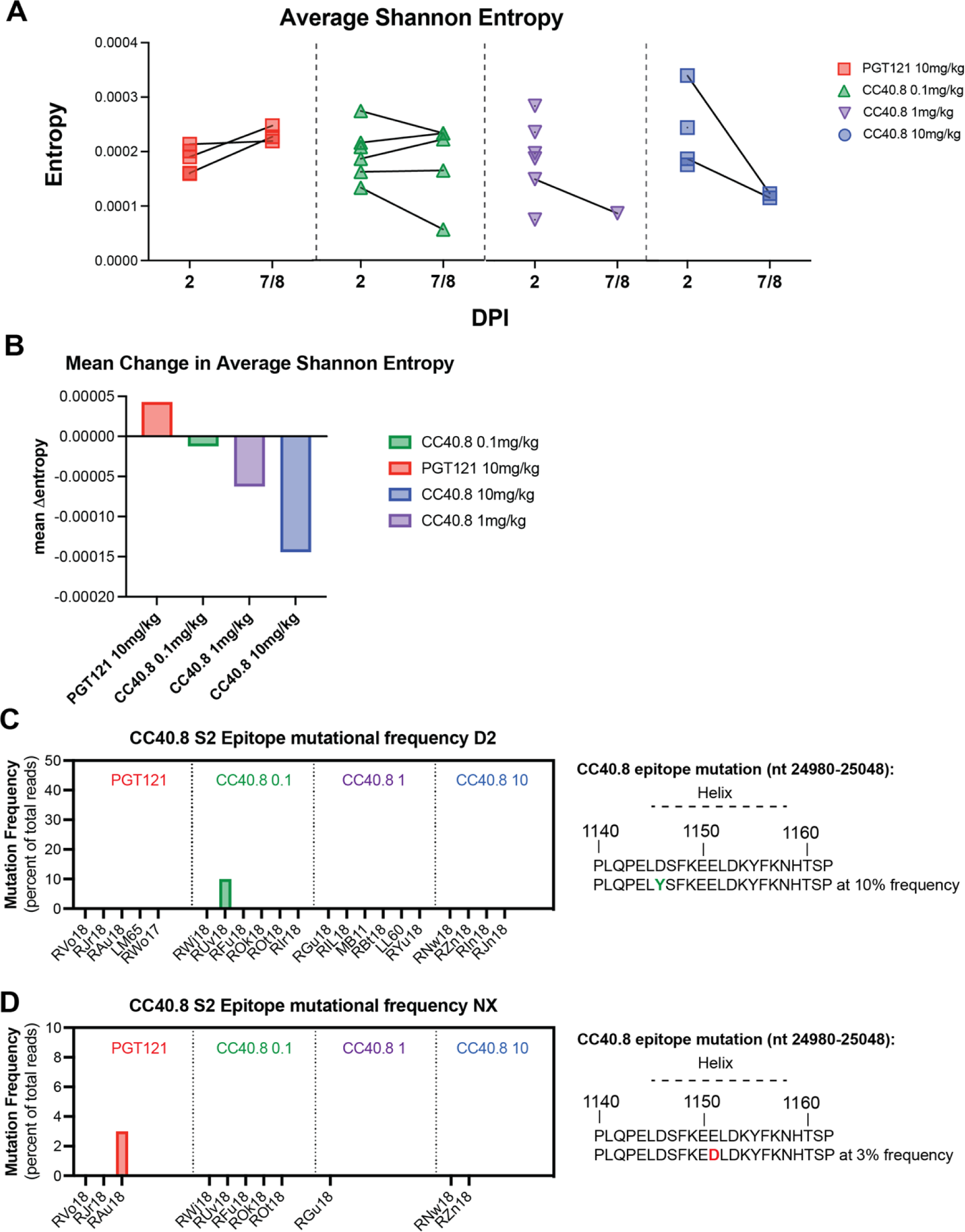
Effect of CC40.8 treatment on SARS-CoV-2 mutant frequency. (**A**) The average Shannon entropy calculated from intra-sample single nucleotide variant (iSNV) frequency in replicate SARS-CoV-2 ARTIC libraries generated from BAL supernatant at 2 dpi and NX (7/8 dpi). Black lines connect libraries from same animal at different timepoints (**B**) Mean change in Shannon entropy for each treatment group from 2 dpi to 7/8 dpi. (**C** to **D**) Frequency of intra-sample, single nucleotide variations at the CC40.8 S2 stem helix epitope at (**C**) 2 dpi and (**D**) 7/8 dpi.

To evaluate the stability of the CC40.8 epitope during SARS-CoV-2 infection, we assessed the frequency of consensus changes and iSNVs within a 23-amino acid segment of the conserved stem helix region within the S2 subunit (nt positions 24980-25048; AA positions 1140-1162). We observed two mutations in the CC40.8 epitope: one animal in the control group exhibited a mutation at E1151D, which was determined to be a contact residue for CC40.8, at 3% frequency at necropsy (Fig 5D). One animal in the 0.1 mg/kg treatment group exhibited mutation D1146Y present at 10% frequency at 2 dpi but was not detected within the samples taken at necropsy (Fig. 5C). These results suggest a lack of antibody escape despite selective pressure, underscoring the conserved nature of the epitope.

## DISCUSSION

Owing to both the ongoing emergence of SARS-CoV-2 variants that circumvent vaccine-elicited immunity, and the zoonotic potential of new CoVs, the development of pan-CoV therapeutic and preventative strategies remains a biomedical priority (70). Reverse vaccinology approaches have identified conserved molecular targets on the coronavirus spike protein capable of eliciting antibodies in humans with broad coronavirus neutralizing capacity, including an epitope contained in the S2 subunit of the coronavirus spike protein (35, 56, 71, 72). Vaccine strategies that elicit neutralizing antibodies by S2 directed binding thus have the potential to reduce the reliance on boosts for novel SARS-CoV-2 VOCs and may provide more protection against novel CoVs. In prior work, we isolated the S2 targeting mAb CC40.8(56), which was capable of neutralizing clade 1b and clade 1a ACE2 receptor-using sarbecoviruses, and had robust *in vivo* protective efficacy against WA.1 SARS-CoV-2 challenge in small animal models (35). A key step in the development of S2 targeting antibodies as a viable strategy for vaccines against coronaviruses is to demonstrate their protective efficacy in a clinically relevant animal model.

Here, we demonstrate that infusion of rhesus macaques with the S2 directed anibody CC4.08 was able to provide protection against SARS-CoV-2 replication in the lower airway. Treatment with mAb CC40.8 resulted in reduced viral load within the lower airway, as well as reduced inflammation following SARS-CoV-2 infection. While we did not observe sterilizing immunity in either compartment, it is worthwhile to note that the inoculation dose, 1.1 x 10^6^ PFU, administered to both the upper and lower airway, is significantly higher than physiological exposure to SARS-CoV-2, yet CC40.8 monotherapy at 1 mg/kg and 10 mg/kg was able to reduce virus by three orders of magnitude. This level of reduction had a protective effect, as RMs with lower viral load exhibited significantly lower levels of infiltrating macrophage populations, expression of ISGs, inflammasome, and other inflammation-associated genes, as well as lower levels of IFNα and other inflammatory mediators detected within the BAL. These data support the working model that links the magnitude of viral loads within the lower airway to the strength of the resulting SARS-CoV-2 disease pathology (61, 66, 69). In this model, higher viral loads within the lower airway result in a more sustained and systemic IFN-I response, which accompanies greater losses of tissue resident alveolar macrophages and greater influxes of macrophage and monocyte subsets that drive the inflammatory milieu (61, 66, 69, 73-75).

In our model, animals with high levels of pre-infused neutralizing antibodies harbored lower levels of SARS-CoV-2 within the lower airway, but did not show a reduction in the upper airway. Reduced availability of CC40.8 within the tissues of the upper compared to lower airway may contribute to the observed differences in viral load reductions. RM in our study were infused with IgG1 subclass mAbs, which lack the J-chain contained with polymeric IgA and IgM isotype antibodies, a polypeptide necessary for trans-epithelial secretion (76, 77). Upper airway viral loads have been associated with transmission risk, and studies have shown a dose-dependent, vaccine-mediated reduction in infectiousness of SARS-CoV-2 breakthrough infections, likely due to reductions in upper airway viral loads (78-85). Recent studies have documented signicant increases in neutralizing IgA titers at the mucosae following intramuscular mRNA vaccination against SARS-CoV-2 (86-89). While as a monoclonal therapy CC40.8 did not reduce upper airway viral loads significantly, a S2 directed antibody response including IgA isotype antibodies, such as those elicited by SARS-CoV-2 vaccination or infection, may prove more effective at reducing upper airway viral loads and transmission risk.

Humoral protection in humans against SARS-CoV-2 infection is conferred by combination of vaccinations, boosters, and prior infections, yielding a vast range of immune states across the globe. There are currently two major contributors to SARS-CoV-2 “breakthrough” infections: waning vaccination or infection-induced immunity, and the evolution of new variants of SARS-CoV-2 with greater immune escape (33, 36, 45, 90-94). Several rounds of boosters have been approved to append the current vaccination series to mitigate the observed waning immunity. Both homologous and heterologous mRNA platform-based boosters have been shown to recover the neutralizing antibody response temporarily against variants of SARS-CoV-2, including Omicron sub-lineages (93, 95-98). Although these mRNA boosters exhibit lower overall vaccine effectiveness at preventing infection against current circulating strains than the original mRNA vaccine demonstrated during earlier phases of the pandemic, they still exhibit remarkable and durable protection against severe disease outcomes. Our data supports maintaining high enough levels of neutralizing antibody through boosters to limit viral replication within the lower airway, thus limiting the magnitude of subsequent lower airway inflammation.

COVID-19 booster doses have employed both the ancestral spike immunogens utilized by the initial vaccination series as well as variant specific spikes, such as the BA.4/BA.5 bivalent booster (99). However these variant-specific constructs have shown marginal superiority to ancestral immunogens at eliciting Omicron-specific protection, possibly due to immunogenic bias from existing humoral immunity away from novel, Omicron-specific epitopes on the spike protein (100). Another concern with designing immunogens to elicit variant-specific responses is, in the time a variant specific immunogen can be tested, produced, and delivered, new variants can displace the dominant circulating strain. A vaccine strategy designed to include a component to elicit neutralizing antibodies directed at the conserved S2 region could circumvent the need for annual updating of COVID-19 boosters to reflect circulating strains (72). Studies have shown individuals with cross-reactive antibodies to endemic HCoVs have higher survival rates from severe disease and protection from infection, further supporting development of a bnAb-targeted CoV immunogen (31, 32, 101).

Our data also provides important preclinical insights surrounding mAb therapies for the treatment and prophylaxis of COVID-19. Based on their loss of neutralization capacity against the Omicron sub-variants, the five previously approved mAb therapies for COVID-19 under emergency use authorizations have all been suspended or revoked by the FDA and no current mAb therapies remain in use(92, 102-109). Our dose-dependent reductions in viral load due to preinfusion with mAb CC40.8, dramatic reductions in inflammation in 10 mg/kg treated animals and the broad reactivity of CC40.8 all suggest that S2-targeting mAb therapies may serve as a preventative strategy in those at high risk of contracting COVID-19 or treatment for those who are infected and at high risk of progressing to severe disease. Additionally, evidence suggests that S1-directed mAb treatment in immunocompromised individuals can promote the emergence of SARS-CoV-2 escape mutations (110). While this study does not attempt to model immunocompromised humans, the lack of S2 mutations observed, coupled with the decreases in mutational entropy in CC40.8-treated animals, warrants further investigation of S2 targeting as a potential treatment strategy for immunocompromised individuals. While frameworks outlining the difficulty of SARS-CoV-2 to maintain selective fitness in humans when mutating the conserved stem helix epitope is largely supported by the conservation of the epitope across β-CoVs, some studies have assessed the positional Shannon entropy of each amino acid position within the SARS-CoV-2 spike protein and identified mutation “hot spots” (111). The CC40.8 epitope lies outside any identified “hotspot,” supporting the conserved nature of the CC40.8 epitope within humans (30).

Although we show that CC40.8 mAb partially protects RM against SARS-CoV-2 infection in this study, there are several limitations. First, SARS-CoV-2 infection in RMs models mild to moderate COVID-19 disease and the extent of protection this S2 stem helix bnAb treatment provides against severe disease in clinical models must still be investigated (57, 61, 65, 69, 74). Second, this study investigated protection against the WA.1 strain of SARS-CoV-2; testing the protective efficacy against more contemporary variants as well as other β-CoVs will be important. Third, this study did not address the contribution of antibody effector functions to protection or, conversely, any mechanisms of antibody dependent enhancement of infection, both warranting further preclinical studies.

In conclusion, this study demonstrates the efficacy of a first-generation mAb, CC40.8, targeting a conserved, cross neutralizing β-CoV epitope at reducing *in vivo* viral replication and mitigating the disease pathology of SARS-CoV-2 infection within the lower airway of a clinically relevant animal model. Since this onset of this study, several mAbs targeting the conserved stem helix epitope of CoVs have been identified with considerably greater neutralization potency and breadth than CC40.8 (112). Overall, this study supports furtherance of experimental and clinical development of S2-targeting antibodies as a strategy to protect and treat coronavirus infection.

## MATERIALS AND METHODS

### Study Overview

An overview of the study design outlined in **Fig. 1a**. 24 RMs were infused intravenously 5 days pre infection with either a 0.1 mg/kg, 1mg/kg, or 10mg/kg concentration of experimental mAb cc40.8 or with control mAb PGT121, with each experimental group consisting of 6 animals (1 female and 5 males). Animals were screened for preexisting, SARS-CoV-2 spike specific antibodies prior to enrollment in this study. Preinfection baseline samples were collected at -4 dpi. At 0 dpi, all RM were inoculated with 1mL intranasally and 1mL intratracheally with a combined total of 1.1 x 10^6^ plaque forming units (PFU) of SARS-CoV-2 (2019-nCoV/USA-WA1/202). BAL and nasal swabs were collected from inoculated animals at 2dpi and at the time of necropsy (7 or 8dpi), with viral titers peaking in these tissues at 2dpi in the infected animals.

#### Sex as a Biological Variable

24 rhesus macaques of Indian origin were sorted by sex, age and weight and then stratified into 4 groups (n=6). Our study examined male and female animals, and similar findings are reported for both sexes.

### Animal Models

Animals used in this study were 24 (6 females and 18 males; mean age of 5 years and 11 months old) specific-pathogen free (SPF) Indian-origin rhesus macaques (RM; Macaca mulatta; Table S1). Animals were initially housed in ENPRC’s BSL2 facilities. Pre-existing, SARS-CoV-2 spike binding antibodies were below detectable levels in all animals prior to infusion. On Day -5, animals in groups of six were infused intravenously with either the control antibody PGT 121 at 10mg/kg or various doses of CC40.8 (10mg/kg, 1mg/kg, 0.1 mg/kg). All antibodies were supplied in solution and diluted with DPBS. Animals were moved to the ABSL3 facilities on Day -4 following baseline collection for a 4 day acclimatization period before infection. One animal was moved on Day -3 due to vomiting during baseline BAL collections on Day -4. On Day 0, animals were inoculated with 1.1 x 10^6^ plaque forming units (PFU) of SARS-CoV-2 (2019-CoV/USA-WA1/2020) via 1mL intratracheally and 1mL intranasally (0.5mL per nostril). After infection, animals were monitored daily by cageside observations which measured responsiveness, discharge, respiratory rate, respiratory effect, cough, and fecal consistency (Supplementary Table S2). In addition, during each anesthetic access, body weight, body condition score, respiratory rate, pulse oximetry, rectal temperature were recorded along with a clinical assessment of discharge, respiratory character and hydration (Supplementary Table S3). Over the 12/13 day period from baseline to necropsy, the following tissues were collected from animals: peripheral blood, bronchoalveolar lavage (BAL) and nasal swabs of both nostrils with the addition of right caudal lung, spleen and hilar lymph nodes at necropsy. Additionally, right middle lung, right caudal lung, right cranial lung, left caudal lung, jejunum and ileum were collected for immunohistochemistry.

### Viral Stock

SARS-CoV-2 WA1/2020 (10/23/21) stock virus with a titer of 3.2 x 10^6^ pfu/mL was provided by the Virus Characterization Isolation Production and Sequencing (VCIPS) Core at Tulane National Primate Research Center. Stock virus was also sequenced to determine the original virus sequence. Using 140uL of the viral stock place in AVL buffer, RNA was extracted using the QiaAmp Mini RNA Viral Kit (#52904). Using 8uL of RNA elution, cDNA is created using the Superscript III First-Strand Synthesis (#18080-051). Next the cDNA is put through NEBNext Ultra II Non directional RNA Second Strand Synthesis Module (NEB Cat #E6111S/L). Finally a PCR clean up is done using the PureLink PCR Purification Kit (#K3100-01/02).

### Determination of viral load RNA

SARS-CoV-2 gRNA N, sgRNA N and sgRNA E were quantified in NP swabs and BALs by 2 independent sites as described in the Supplementary Materials.

### Tissue SARS-CoV-2 RNA Quantification

Lung tissue was harvested on 7 or 8 dpi and homogenized using Bead Ruptor 12 (Omni International). Modified protocol from Zhou et al. 2022(30).

### Expression and purification of monoclonal antibodies CC40.8 and PGT121

mAbs CC40.8 and PGT121 were produced as described in the Supplementary Materials. Modified from Zhou et al. 2023(41).

### Anti-Spike Antibody Detection in BAL Supernatant and Serum Samples by ELISA

Anti-spike ELISA’s were performed asdescribed in the Supplementary Materials.

### Tissue collection and processing

PB, NP swabs, throat swabs, and BAL were collected longitudinally. At necropsy, lower (caudal) lung, upper (cranial) lung, and hilar LNs were also collected. Detailed methods pertaining to the collection and processing of these tissues are included in the Supplementary Materials

### Single-cell RNA-Seq Library and sequencing from NHP BALs and Lung

Single cell suspensions were prepared and loaded onto the 10X Genomics Chromium Controller in the BSL3 facility using the Chromium NextGEM Single Cell 5’ Library & Gel Bead kit to capture individual cells and barcoded gel beads within droplets (113). The libraries were prepared according to manufacturer instructions, including the preparation of feature barcode libraries for hashtag detection. They were then sequenced on an Illumina NovaSeq 6000 with a paired-end 26x91 configuration targeting a depth of 50,000 reads per cell. Cell Ranger software was used to perform demultiplexing of cellular transcript data, as well as mapping and annotation of UMIs and transcripts for downstream data analysis.

### Single-cell RNA-Seq bioinformatic analysis of BAL and Lung cells

The cellranger v6.1.0 (10X Genomics) pipeline was used for processing the 10X sequencing data and the downstream analysis was performed using the Seurat v4.0.4 R package. A composite reference comprising of Mmul10 from Ensembl release 100 and SARS-CoV2 (strain MT246667.1 - NCBI) was used for alignment with cellranger. The percentage of SARS-CoV-2 reads was determined using the PercentageFeatureSet for SARS-CoV2 genes. For BAL samples, a total of 107,830 cells across all animals passed quality control (QC) and were used for analyses. For lung samples, a total of 101,766 cells passed upstream QC and were used for analysis. The bioinformatic processing of scRNA-Seq data and subsequent analysis was performed as described previously for BAL samples (66) and lung samples(69). Detailed methods pertaining to the bioinformatic analysis are included in the Supplementary Materials

### SARS-CoV-2 ARTIC Library Generation

SARS-CoV-2 ARTIC cDNA libraries were generated from RNA recovered from BAL supernatant at 2 dpi and at NX. Detailed methods pertaining to the generation of these libraries are included in the Supplementary Materials

### SARS-CoV-2 Sequence Analysis

SARS-CoV-2 reference-based assembly was performed with nf-core/viralrecon v2.6, using default parameters with no trim offset (114, 115). First, a consensus sequence was assembled from the reads generated from the infecting virus sample (using reference sequence MN908947.3), and this full-length SARS-CoV-2 sequence (“Input_consensus”) was subsequently used at the reference sequence for assembly and variant calling from the reads generated from each experimental sample. Only samples with at least 95% genome coverage in both replicate libraries were included in sequence analysis (Supplemental Table X). These samples had a median depth of coverage across the genome ranging from 25,772X to 119,292X (median 48,563X). Intra-sample single nucleotide variants (iSNVs) were called against the “Input_consensus” sequence with iVar v1.3.1, setting the maximum depth at 29 million bases, minimum quality score at 15 and minimum frequency at 1% (116). Further filtering was used to identify iSNVs present in two replicate libraries, and at positions with at least 100X depth. Using the average frequency of each iSNV from the two replicate libraries, average Shannon entropy for each sample was calculated as the sum of (-ln(frequency)*frequency)/29903, where frequency indicates the allele frequency of each iSNV, and 29903 is the length of the reference genome sequence.

### Macrophage Flow Cytometry Immunophenotyping

Macrophage immunophenotyping was performed as described in the supplementary materials.

### Statistical analysis

All statistical analyses were performed two-sided with P ≤ 0.05 deemed significant. Ranges of significance were graphically annotated as follows: *P < 0.05; **P < 0.01; ***P < 0.001; ****P < 0.0001. Analyses for Figs. 1 (B and C), 2 (B to G), 3 (C to F, K to M), and figs. S1 (A to L), S3 (B to E), S4 (A to J), S6 (D to G) were performed with Prism version 10 (GraphPad).

### Study Approval

Emory’s National Primate Research Center (ENPRC) is certified by the U.S. Department of Agriculture (USDA) and by the Association for Assessment and Accreditation of Laboratory Animal Care (AAALAC). All animal procedures were completed in line with institutional regulations and guidelines set by the NIH’s Guide for the Care and Use of Laboratory Animals, 8th edition, and were conducted under anesthesia with appropriate follow-up pain management to ensure minimal animal suffering. All animal experimentation was reviewed and approved by Emory University’s Institutional Animal Care and Use Committee (IACUC) under permit PROTO202200025 and reuse

## Data and Materials Availability

Data tables for expression counts for single-cell RNA-seq for BAL are deposited in NCBI’s Gene Expression Omnibus and are accessible through the Gene Expression Omnibus (GEO) under accession number (Pending). The processed single-cell lung macrophage reference dataset (66) was originally obtained from GEO under accession no. GSE14975833 (67). Custom scripts and supporting documentation on the RNA-seq analyses will be made available at https://github.com/BosingerLab/NHP/COVID_mAb Reagents generated in this study may be requested from the authors with a completed materials transfer agreement. All data needed to evaluate the conclusions in the paper are present in the paper or the Supplementary Materials.

## AUTHOR CONTRIBUTIONS

CTE, KAK, EG, RA, TR, DRB and SEB conceptualized the study. RA identified and characterized CC40.8. EG produced and validated safety of CC40.8 antibody stocks. SAL wrote the IACUC protocol for the animal studies. SAL, TT, and MCL processed all RM blood samples in an ABSL-2 facility. MP provided critical input in the development of tissues collection protocols and alveolar macrophage flow cytometry methodology. CTE, KAK, and EG processed all SARS-CoV-2–infected samples in an ABSL-3 suite with assistance from HA. CTE performed MSD analysis on BAL fluid from uninfected and SARS-CoV-2– infected RMs with assistance from TT and MCL. MR, SW, JSW, and AW conducted all longitudinal animal collection procedures for SARS-CoV-2–infected RMs in the ABSL-3 and EHC performed necropsy collections. MG, CCH and MRB performed repeat measurements of sgRNA-N and sgRNA-E viral loads in nasopharyngeal swab and BAL. NG performed repeat measurements of sgRNA-E, sgRNA-N, and gRNA-N viral loads in nasopharyngeal swabs and BAL. CTE, KAK, and TT performed multiparameter flow cytometry and CTE analyzed flow data. EG assayed lung tissue viral loads. EG assayed serum-neutralizing antibody titers and NB analyzed BAL antibody titers. H.A. performed 10X Genomics scRNA-seq, and CTE performed preprocessing for single-cell BAL data and conducted and graphed all scRNA-seq analyses with oversight from AA. AM and SAL prepared SARS-CoV-2 ARTIC libraries from BAL supernatant. Mutational entropy analysis was performed by AB and AP. Funding was acquired by DRB (supplement to UM1AI44462 and by the Bill and Melinda Gates Foundation (INV004923). CTE and SEB wrote the manuscript with KAK, EG, DRB, RA and TFR providing critical input. Order of first authors was determined by amount of contribution towards writing the final manuscript.

## ACKNOWLEDGMENTS

We thank the Emory National Primate Research Center (ENPRC) Division of Animal Resources for providing support in animal care. We thank Kalpana Patel from the Environmental Health and Safety Office for BSL-3 training and safety oversight. We thank Dr. Elise Viox and Diane Carnathan for their input in study planning and preparation. We thank Dr. Mehul Suthar for his input on SARS-CoV-2 stock selection. We thank Jianjun Chang at the Emory Multiplexed Immunoassay Core for help with MSD assay. We would like to thank the High Containment Research Performance Core: RRID: SCR_024612; at Tulane National Primate Research Center for technical assistance.

## LIST OF SUPPLEMENTARY MATERIALS

Fig. S1 to S9

Tables S1 to S4

Data Files S1-S8

MDAR Reproducibility Checklist

References (*117-129*)

